# Long-Term Potentiation and Closed-Loop Learning in Paired Brain Organoids for CNS Drug Discovery

**DOI:** 10.1101/2025.07.03.663054

**Authors:** Corey Rountree, Eva Schmidt, Nicholas Coungeris, Victoria Alstat, Andrew S. LaCroix, Megan Morris, Michael J. Moore, J. Lowry Curley

## Abstract

Learning and memory, central to cognitive function and critical targets for drug discovery in neurological disorders, fundamentally rely on synaptic plasticity, such as long-term potentiation (LTP). We developed an embedded electrode array (EEA) capable of supporting paired CNS-3D brain organoids, derived from human, induced pluripotent stem cells (iPSCs), that were interconnected via guided axonal growth, as a novel functional biomarker model for probing learning-related plasticity *in vitro*. Using open-loop stimulation protocols, a robust, network-level LTP in trained organoids was demonstrated that was not observed in untrained organoids. Importantly, LTP was enhanced in the presence of elevated levels of brain-derived neurotrophic factor (BDNF), and it was prevented by the NMDA receptor antagonist AP5. Furthermore, a closed-loop “maze-game”, inspired by Pac-Man®, demonstrated that paired brain organoids could learn to perform better in the game when given reinforcement feedback, and that this effect depended on BDNF. This integrated platform provides a human-relevant *in vitro* model of synaptic plasticity and cognitive function, establishing a functional biomarker platform for central nervous system (CNS) drug discovery and linking game performance as a measurable readout of learning-related plasticity to the emerging field of organoid intelligence.

## Introduction

Developing accurate models of human learning and memory is essential for accelerating drug discovery targeting devastating neurological disorders, including Alzheimer’s and Parkinson’s diseases. Despite intensive efforts, therapeutic development for central nervous system (CNS) diseases have high failure rates (∼100%), largely due to models and biomarkers that do not accurately predict human efficacy^1–4^. For instance, currently available biomarkers for Alzheimer’s disease include amyloid-beta plaques and tau tangles, which can be detectable years before symptom onset but are most prominent and clearly diagnostic in later disease stages^5^. Although these biomarkers have significantly advanced understanding of disease progression, they primarily reflect downstream pathological features rather than directly indicating functional disruptions that precede cognitive decline^6,7^. Moreover, these pathological features have proven difficult to replicate in traditional cellular and animal models, contributing to the high failure rates for CNS-focused drugs^1^. Consequently, in order to accelerate the discovery of effective CNS drugs, there is an urgent need for more functional, human-relevant biomarkers that can robustly predict therapeutic outcomes.

Central to learning and memory are synaptic plasticity mechanisms, particularly long-term potentiation (LTP), which involve activity-dependent strengthening of synaptic connections over the course of hours to days^8,9^. Dysfunction in these processes is closely linked to cognitive impairments seen in diseases such as Alzheimer’s and schizophrenia^10,11^. Although extensively characterized in animal models and isolated human brain slices, investigating LTP directly in human tissues at the scale for drug discovery remains challenging due to limited tissue availability^12,13^. Recent work, however, suggests that these mechanisms may be modeled in human-derived *in vitro* systems^14–17^, creating an opportunity to develop more functional, human-relevant biomarkers for CNS therapeutics. Despite this promise, dissociated iPSC-derived networks often lack the multicellular architecture, maturation, and long-range connectivity that shape neural circuits in the human brain, motivating the use of human brain organoid models^18–21^.

Three-dimensional (3D) human brain organoids derived from induced pluripotent stem cells (iPSCs) have emerged as a promising platform to meet this need^22^. These tissues develop diverse neuronal and glial populations^23–25^, exhibit spontaneous oscillatory activity reminiscent of neonatal EEG^26,27^, and can model a broad range of neurological diseases^28–35^. They also express key molecular components of synaptic plasticity,^15,36,37^ although robust and persistent network-wide LTP has not been clearly established. One likely limitation is that conventional organoid cultures lack the structured long-range connectivity required for organized circuit function. In response, engineered multi-organoid systems such as assembloids, in which organoids are fused together, and “connectoids,” in which organoids are cultured at a distance to allow axonal connections to connect them, have begun to provide more physiologically relevant inter-organoid communication and improved functional maturity^37–41^. Such platforms also intersect with the emerging field of organoid intelligence (OI), in which electrophysiological stimulation and readout are used to interrogate adaptive, learning-relevant responses in human neural tissues^42–46^. These advances position brain organoids as promising human-relevant models for studying synaptic plasticity and accelerating CNS drug discovery^21,27,47^.

Building on these insights, we developed a 24-well embedded electrode array (EEA) platform that supports paired CNS-3D brain organoids with guided axonal connectivity across extended electrode channels. This structured system enables chronic, spatial electrophysiological probing of both spontaneous and evoked neural activity. Using learning-inspired stimulation protocols, we demonstrated robust, pharmacologically modifiable LTP in organoid networks. In a closed-loop maze paradigm inspired by Pac-Man® and reinforcement learning principles, organoids improved their game performance in response to reward- and penalty signals, consistent with functional plasticity and rudimentary learning. These findings establish the organoid-EEA platform as a versatile in vitro model for studying learning-related plasticity, with potential applications in CNS drug discovery.

## Results

### Paired CNS-3D organoids establish axonal connections across millimeter distances

Upon placement in the EEA co-culture plate^48^, organoid pairs readily extended axons into the connecting channel directed towards one another, achieving physical and functional integration within 5 weeks. Figure 1A shows an immunofluorescence image of a representative day 35 CNS-3D cortical brain organoid^27,49^ stained for MAP2 (neurons, green) and GFAP (astrocytes, magenta), illustrating the cell composition and size (∼500 µm) of the organoids. The custom 24-well EEA plate (Fig. 1B) allowed two organoids to be positioned in each well with a defined separation of 2-9 mm. A narrow polyimide microchannel (200 µm width, 500 µm height) guided neurites to grow directly toward one another across the intervening gap, wherein 10 planar microelectrodes were embedded in the channel floor to record and stimulate the growing axon bundle. We empirically tested three different inter-organoid distances (2 mm, 4 mm, 9 mm) to optimize network formation. Shorter separations promoted faster physical contact between organoids (∼1-2 weeks) while longer distances required up to 5 weeks for axon bridging. In practice, all configurations eventually yielded a continuous axon tract linking the organoids, typically within 2–5 weeks of co-culture. The formation of these “wired” organoid pairs was highly reproducible across 408 organoid pairs (96 pairs at 2 mm, 120 at 4 mm, 192 at 9 mm). Importantly, the organoid-axon-organoid assemblies remained viable and exhibited oscillatory spontaneous activity and electrically-evoked responses for at least 10 weeks (Fig. 1C), confirming functional integration.

**Figure 1.**
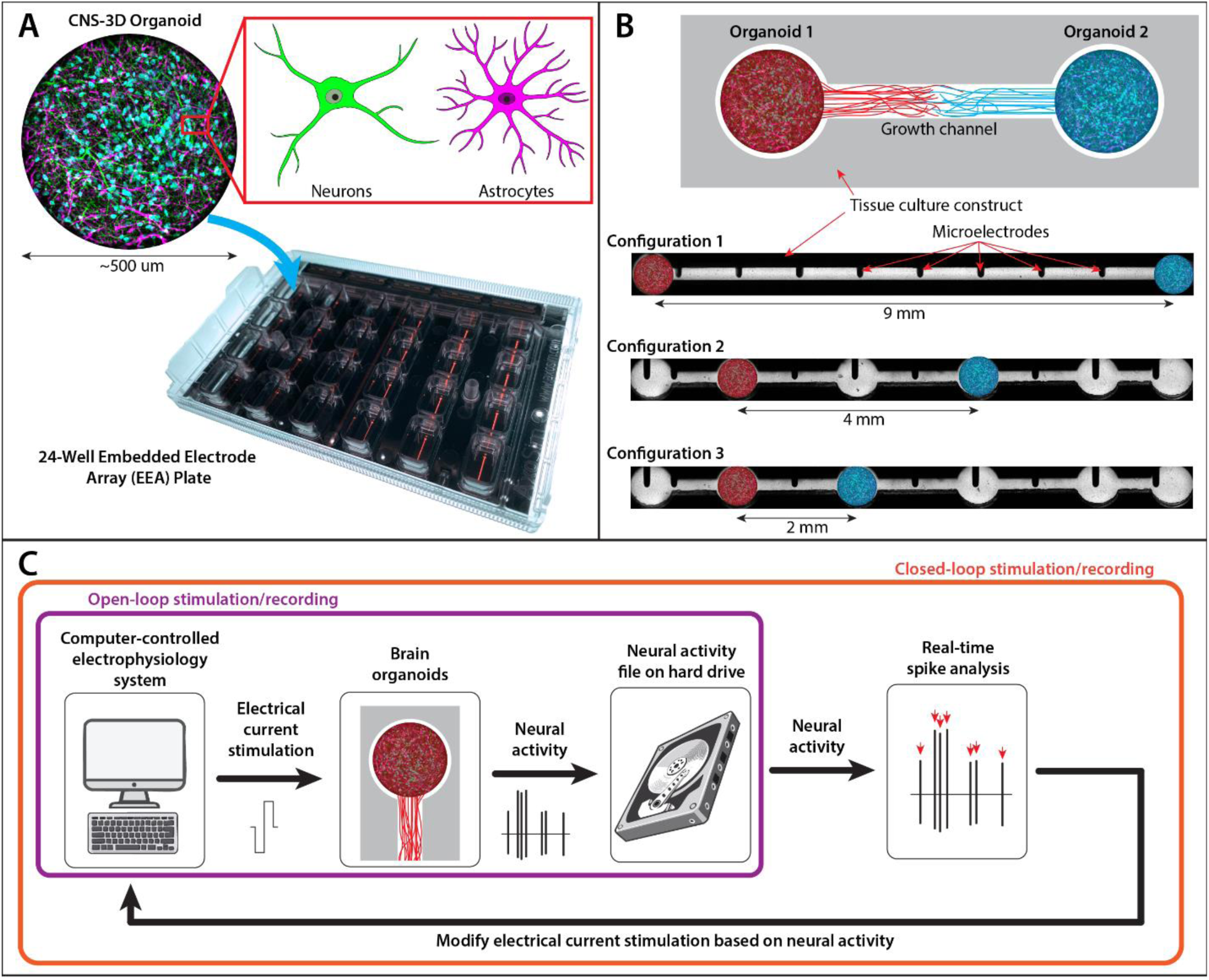
CNS-3D brain organoids connected across a microelectrode plate. **A)** Immunofluorescence micrograph of a human iPSC-derived CNS-3D brain organoid (∼500 µm Ø) composed predominantly of cortical neurons and astrocytes (representative markers: MAP2 [green] & GFAP [magenta]). **B)** Schematic of the custom 24-well embedded-electrode array (EEA) plate used for dual-organoid co-culture. Two organoids are positioned in each well and linked by a growth channel that guides axonal projections across ten planar microelectrodes embedded in the channel floor. Three inter-organoid separations were tested: Configuration 1 = 9 mm, Configuration 2 = 4 mm, and Configuration 3 = 2 mm. **C)** Block diagram of open-loop (purple) versus closed-loop (orange) stimulation paradigms. Open-loop delivers predefined current pulses to electrodes and records the evoked neural activity for offline analysis. Closed-loop performs real-time spike detection on recorded signals and dynamically adjusts subsequent stimulation parameters based on the detected activity.

### Physical organoid–organoid connections completed in <5 weeks

Time-lapse brightfield imaging and post hoc immunostaining revealed the spatiotemporal progression of axon growth through the EEA channel (Fig. 2). Figure 2A presents a montage of brightfield images from a representative organoid pair with 9 mm separation over its first five weeks in co-culture (day *in vitro*, DIV 3 to DIV 35). In the initial days, individual neurites emerged from each organoid into the proximal end of the channel. By one week (DIV 7), there was a dense web of fibers near each organoid, evident as bundle-like structures extending into the channel (Fig. 2A, DIV 7, red inset). From week 2 onward, these axonal fronts continued to elongate and eventually converged at the mid-channel (orange inset), establishing physical contact between organoids with increasing fiber density through DIV 35. By DIV 35, the phase-contrast image shows an obvious tissue strand spanning the entire channel. Correspondingly, immunofluorescence labeling for β-III tubulin at 9 weeks to highlight axonal fibers (Fig. 2B) confirmed a continuous, fasciculated axon bundle bridging Organoid 1 and Organoid 2 (insets highlight the tightly bundled tract at each organoid and in the mid-channel). Axons were observed to intermingle within this bundle, creating a potential feed-forward and feedback loop between the two organoids.

**Figure 2.**
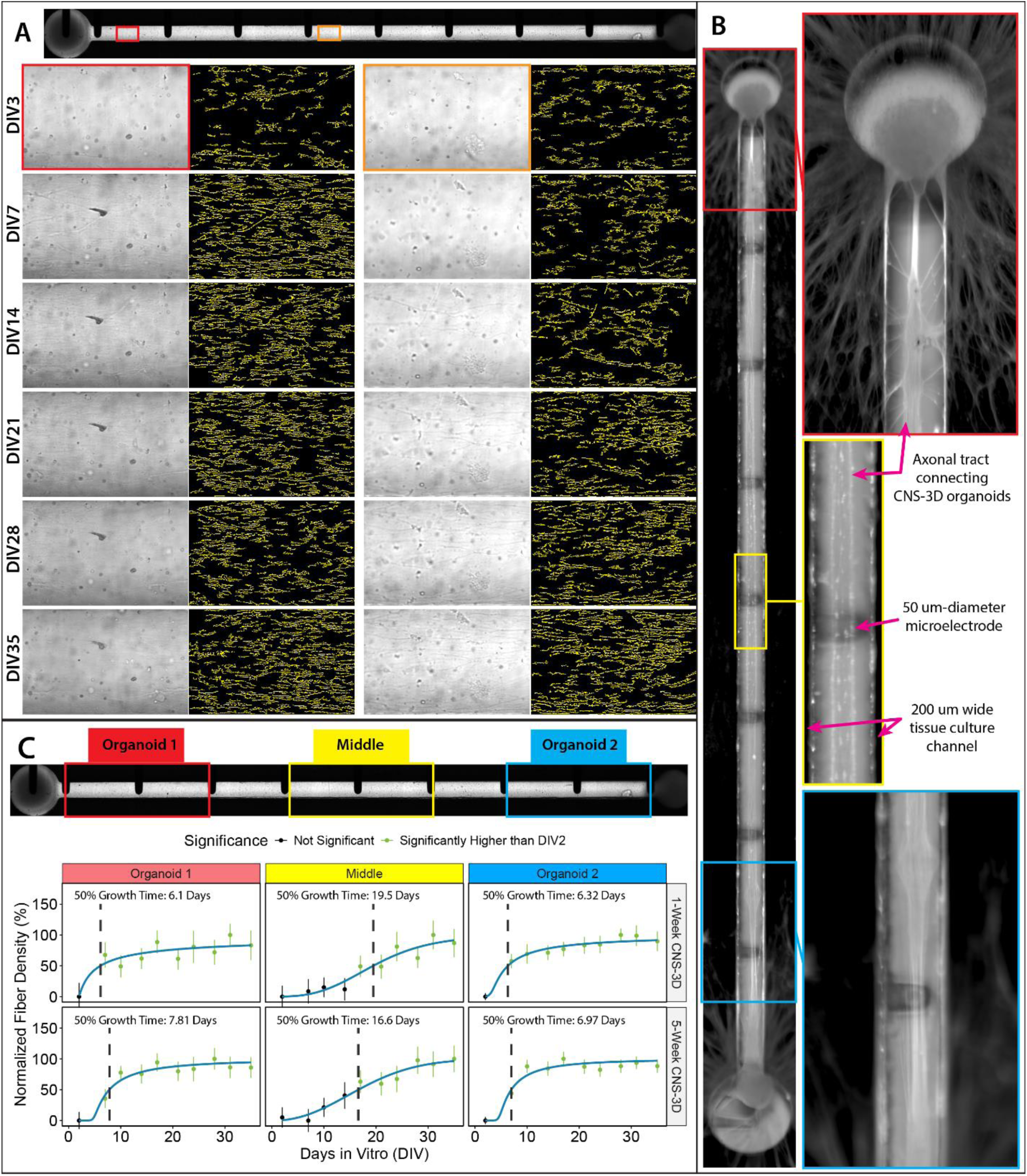
Physical connections between organoids occurred within 2-3 weeks. **A)** Brightfield montage of a representative 1-week-old CNS-3D organoid pair (day in vitro, DIV 3–35). The red and orange boxes mark regions adjacent to Organoid 1 and the channel midpoint, respectively. For each region, brightfield closeups (left) and niBlack-filtered fiber masks (right) illustrate rapid fiber densification near the organoid by DIV 7, followed by progressive infill of the mid-channel through DIV 35. **B)** Whole-channel immunofluorescence (β-III-tubulin) highlighting Organoid 1 (red), channel midpoint (yellow) and Organoid 2 (blue). Insets reveal a continuous, fasciculated axon bundle bridging the two organoids. **C)** Quantitative fiber-density trajectories (mean ± SEM, normalized to maximum) for the three regions. Top: 1-week organoids; bottom: 5-week organoids. Sigmoid fits (blue lines) yield EC₅₀ values (time to 50 % max density; black dashes) showing regions near the organoids reach max density about 2-3 times faster than the middle of the channel. These data show that axons extend rapidly from each organoid and converge to form a robust, bidirectional nerve bundle whose maturation rate depends on both organoid age and channel position.

To quantify outgrowth kinetics, we measured fiber density over time in three regions: near Organoid 1 (red zone), the channel midpoint (yellow), and near Organoid 2 (blue). Figure 2C plots the normalized fiber density versus time for young organoids (1 week old at placement in the EEA plates; N = 24 organoid pairs) and more mature organoids (5 weeks old at placement in the EEA plates; N = 48 organoid pairs). In all cases, axonal extension followed a sigmoidal growth curve with an initial, rapid phase that plateaued as the channel filled. As expected, fibers reached 50% of maximal density significantly sooner in regions closer to the organoids than in the mid-channel. The half-maximal outgrowth times (EC₅₀) in the organoid-adjacent regions were about 2–3 times faster than at the channel center, reflecting that axons sprouted quickly from the organoid surface and then gradually populated the distal gap. By the end of week 5, organoid-adjacent regions achieved a higher overall fiber density compared to the mid-channel, especially for the older organoid cohort. Nevertheless, both young and mature organoids ultimately formed robust inter-organoid tracts. Together, these data demonstrated that two independently grown organoids can be linked via a self-assembled axonal bundle. The maturation rate of this “mini-circuit” depended on both the developmental stage of the organoids and the distance between them. However, relatively immature organoids also projected long axons and established a connected bridge within a few weeks that became more dense over time.

### Functionally-relevant network activity at 2-3 weeks

Having established structural connectivity, we examined how functional activity in the organoid networks evolved longitudinally with extended culture timelines. Using the embedded electrodes, spontaneous electrophysiological activity was recorded from organoid pairs over multiple weeks with a custom electrophysiology system (Figure 3A). Figure 3B tracks the development of network bursts in a representative organoid pair from 1 to 3 weeks *in vitro* (DIV 6–20). Kernel density estimates of burst frequency are plotted for an electrode beneath one organoid (green) versus an electrode at the channel midpoint (blue). Early on (around 1 week), the organoid showed intermittent, small bursts, while the mid-channel region was mostly silent (no established connection yet). By DIV 13 (∼2 weeks), a clear change occurred: high-amplitude bursting was observed under the organoid with increased regularity, and bursts also appeared on the mid-channel electrode. These phenomena coincided with the initial detection of axonal connection between organoids. By DIV 20 (∼3 weeks), both the organoid and midpoint electrodes exhibited robust network activity arising from one or both organoids. These results align with previous reports that inter-organoid connections amplify network dynamics. In our experiments, organoid pairs consistently showed a marked increase in burst intensity and frequency once the axon bundle formed.

**Figure 3.**
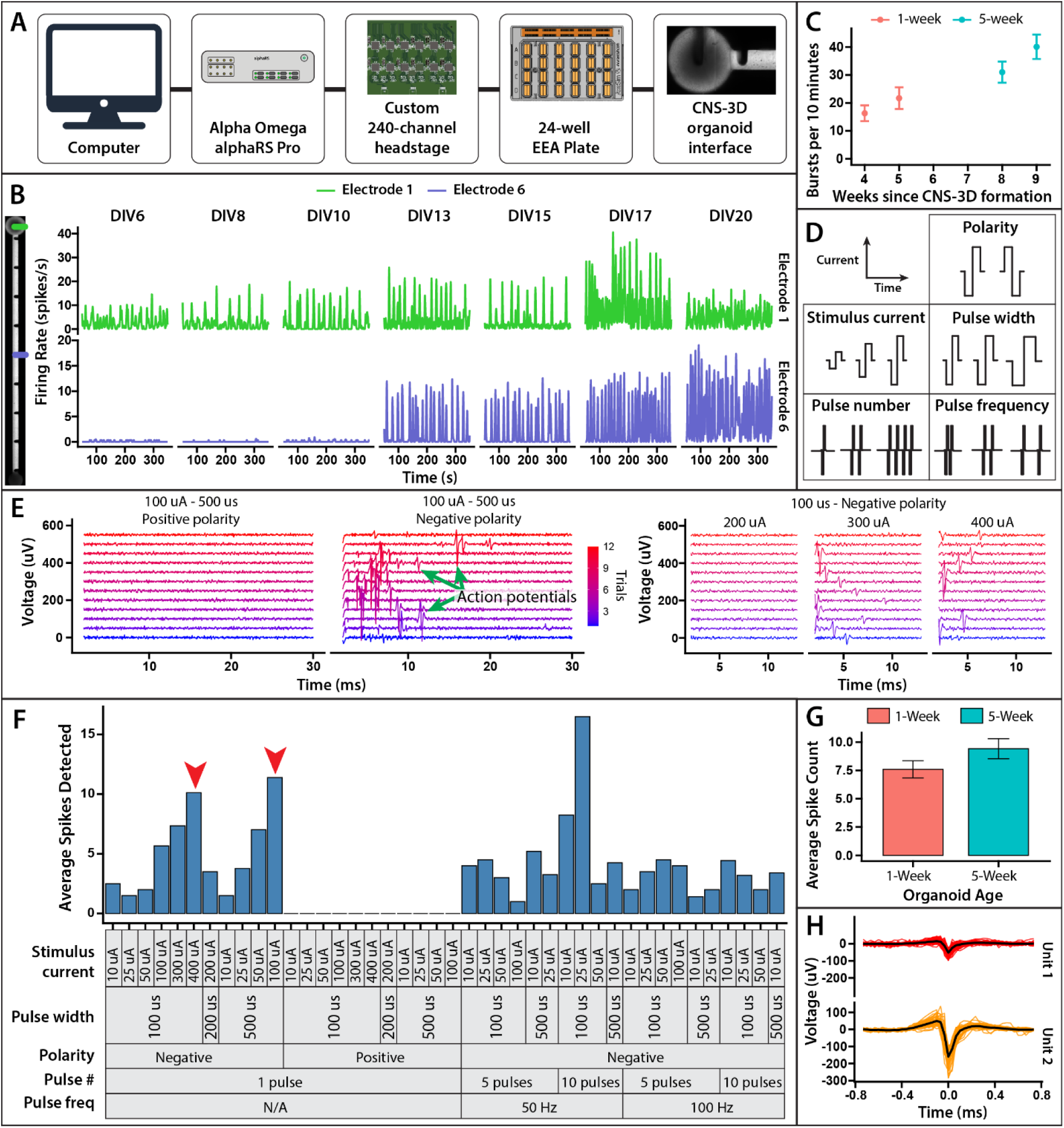
Electrophysiological connections between organoids coincided with physical connections, and 3 effective, non-toxic stimuli were chosen empirically. **A)** Hardware chain for closed-loop electrophysiology. A control PC drives an Alpha-Omega αRS-Pro stimulator/recorder, a custom 240-channel headstage, and a 24-well embedded electrode array (EEA) plate; each well contains two CNS-3D organoids positioned over ten planar electrodes. **B)** Development of spontaneous network bursts. Kernel-density estimates of burst rate recorded from an electrode beneath an organoid (green) and a midway channel electrode (blue) in a representative 5-week culture. Bursting beneath the organoid increases in amplitude and regularity from DIV 6-20, whereas mid-channel bursts appear at DIV 13, reflecting progressive axonal innervation. **C)** Cohort-level maturation. Mean ± SEM number of bursts detected during 10-min recordings versus organoid age (weeks since formation) for cohorts seeded at 1-week (red) and 5-weeks (blue). Trends parallel previously reported FLIPR Ca²⁺ activity of CNS-3D organoids at matched ages. **D)** Parameter space explored for electrical stimulation included: stimulus current, pulse width, initial polarity, pulse number, and pulse frequency. **E)** Representative evoked responses. Overlays of 12 identical trials illustrate a non-response (upper left), a robust multi-unit response (upper right), and a current-ramp series (bottom) showing graded recruitment with increasing current. Trials are vertically offset for clarity with different colors representing early (blue) vs late (red) trials. **F)** Screening outcome. Bar graph summarizes evoked spike counts for every parameter combination (table beneath). Two conditions, 100 µs + ≥300 µA and 500 µs + 100 µA, produced the most consistent, time-locked responses (highlighted) and were selected for all subsequent experiments. **G)** Evoked responsiveness across development. Average spikes ± SEM elicited by the 500 µs, 100 µA paradigm show comparable magnitudes in the 1-week and 5-week cultures (n.s., two-tailed t-test). Therefore, data were pooled for later analyses. **H)** Example spike sorting results following wavelet clustering yields two single-unit waveforms (Unit 1, Unit 2) that were tracked across all subsequent analyses. Together, these results establish reliable metrics for tracking spontaneous network maturation and identify stimulation parameters that evoke robust, stereotyped activity in CNS-3D organoid pairs.

We quantified burst rates across multiple organoid pairs and related those rates to organoid age and initial maturity. Figure 3C summarizes the mean number of bursts per 10 min recording as a function of time in culture, comparing organoids that were seeded into the EEA plates at 1-week-old vs. 5-weeks-old (N = 10, N = 32, respectively). Both cohorts exhibited an upward trend in burst frequency over weeks of co-culture, reflecting ongoing network maturation. The overall activity was higher in the cohort that started with 5-week-old organoids, as expected given their more advanced neuronal development. These trends are consistent with prior calcium imaging studies showing similar bursting rates of CNS-3D organoids at similar time scales. Thus, within our platform, organoid pairs achieved self-organized oscillatory activity that intensified with age, and the presence of inter-organoid connections did not impede the intrinsic maturation of bursting.

### Three non-toxic stimulus conditions determined empirically

We next sought to reliably evoke electrophysiological responses from the organoid networks to probe their functional connectivity and excitability. Using the stimulation paradigm detailed in the Methods, a systematic probing of stimulation parameters was conducted to identify those that best elicited reproducible neural responses (Fig. 3D). The parameter space included current amplitude, pulse width, polarity, pulse number, and frequency. Each condition was evaluated for its capacity to elicit time-locked action potential responses. Figure 3E shows example evoked responses under different conditions. In some cases, a given stimulus produced no response (Fig. 3E, left), indicating that the stimulated site was not closely integrated with the neuronal network or that the parameter set was not appropriate. In other cases, a single pulse triggered a robust burst of spikes (Fig. 3E, middle), often involving multiple units firing within a few milliseconds after the stimulus, which was a sign of functional recruitment of a local microcircuit. We also observed graded responses when varying stimulus intensity: the rightmost panel of Fig. 3E overlays responses to a series of pulses with increasing current (200 to 400 uA), showing incremental spike recruitment with more spikes as current rises. These profiles were reminiscent of input-output curves in traditional brain slice experiments, in which stronger stimulation aggregated more responsive neurons^50^. After systematically exploring stimulus parameters (Fig. 3D), the conditions yielding the most consistent, time-locked activation of the organoid pairs were identified. Figure 3F summarizes the outcome of this screening. Among dozens of tested parameter combinations (pulse widths, amplitudes, etc.), two conditions stood out (red arrows) that evoked reproducible spiking on >80% of trials. These conditions were selected for subsequent experiments: a short duration, high current pulse (100 us, 400 uA; 40 nC/phase) and a long duration, low current pulse (500 us, 100 uA; 50 nC/phase). Repeated stimulation under these conditions over multiple weeks continued to elicit neural activity without detectable changes in imaging, suggesting no appreciable neurotoxicity.

Using these stimuli, we compared the evoked responsiveness of younger vs. older organoid networks. It was found that organoid age did not strongly influence evoked spike yield under our conditions. As shown in Figure 3H, the average number of spikes evoked per stimulus was comparable (p > 0.05) between networks that had been placed at 1-week-old and those placed at 5-weeks-old, with both ages tested after 5 weeks of growth in the EEA plates. This suggests that by the time of these assays (typically 6–10 weeks of total organoid age), both younger-start and older-start networks achieve a similar level of functional connectivity and excitability in response to external input. We therefore pooled data across ages for subsequent plasticity experiments.

We finally performed spike sorting on recorded multi-unit data to assess the complexity and stability of neural signals across the organoid–channel network. As shown in Figure 3H, wavelet clustering of waveforms from a single electrode revealed multiple distinct clusters, each corresponding to a presumptive single unit^51^. This pattern was observed across several electrodes in the system, indicating that multiple individual neurons were recorded at once. The ability to resolve and track multiple units supports the conclusion that our platform samples activity from a distributed population of neurons, consistent with the formation of a functional neuronal network across the paired organoids. Together, results from Fig. 3 establish that our organoid pairs generated reliable metrics of network activity, both spontaneous (bursting) and evoked (stimulus responses). These metrics established a baseline against which we could detect any training-induced potentiation or change in synaptic function.

### Successful induction of LTP that was sensitive to BDNF and NMDA-receptor blockade

To evaluate whether stimulation induced long-lasting changes in network excitability consistent with LTP, we quantified changes in the evoked spike counts from day-to-day evaluations. For this, we implemented an open-loop training paradigm over the course of one week, as described in the Methods. Each day, trained organoids received a series of stimuli (five rounds of pulsing all electrodes), whereas untrained controls did not, while both were evaluated before and after training to measure any changes in evoked spiking. Figure 4A shows spike plots from two representative wells from untrained (top) and trained (bottom) groups recorded during the first evaluation each day, illustrating increased evoked responses across successive days in the trained organoid pair but not the untrained pair. At the population level, LTP was quantified as the ratio of evoked spikes on each day relative to the Day 0 baseline for trained versus untrained groups (N = 19 organoid pairs per condition). Figure 4B shows the mean normalized spike count trajectories over the training week for three conditions: control medium, BDNF-supplemented, and AP5-treated (N = 12 organoid pairs per condition). In control conditions, the trained organoids exhibited a significant increase in evoked spikes compared to untrained by Day 2, and this potentiation became larger by Day 3–4 (trained reaching ∼1.9-fold baseline vs. untrained ∼0.8-fold). The separation between trained and untrained was significant on Days 2–4 (p < 0.05; Table 1) and persisted through the end of the week. Notably, when elevated BDNF was added (50 ng/mL), the rate and extent of potentiation were even greater: trained organoids in BDNF climbed to ∼3-4× baseline by Day 4, nearly double the level of the trained control group. This suggests that exogenous BDNF provided an additional boost, possibly by lowering the threshold or enhancing the magnitude of synaptic strengthening events. In contrast, NMDA receptor blockade with AP5 completely prevented any training effect - the trained and untrained curves under AP5 were indistinguishable throughout the week (both around ∼1.1–1.5× baseline, with no significant difference), consistent with NMDA receptor activity being required for the induction of sustained LTP. These divergent outcomes highlight that the potentiation we observed after extensive training corresponds to long-term potentiation, as opposed to nonspecific increases, given its dependency on NMDA receptor-mediated mechanisms.

**Figure 4.**
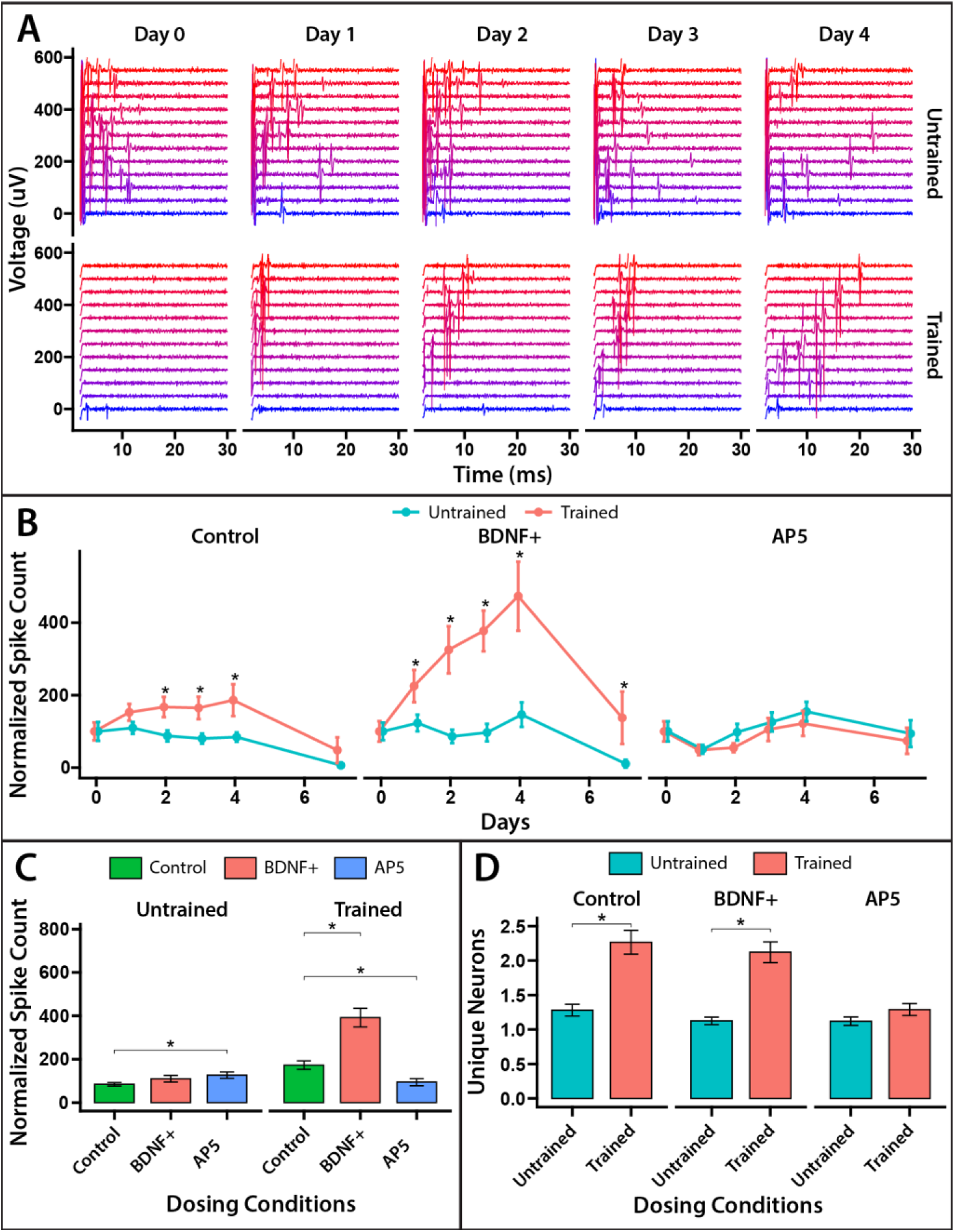
Open-loop stimulation induces LTP that is sensitive to BDNF and NMDA-receptor blockade. **A)** Spike plots from untrained (top) and trained (bottom) wells illustrate the learning-week trajectory: repeated open-loop stimulation during Evaluation #1 drives a day-by-day increase in evoked network activity in the trained, but not the untrained wells, signaling progressive potentiation. **B)** Population data, expressed as spike counts normalized to Day 0 (mean ± SEM), reveal that the control-medium samples display a significant trained-over-untrained elevation from Day 2 through Day 4. Cultures exposed to 50 ng mL⁻¹ BDNF+ potentiate even more rapidly, reaching almost twice the control gain by Day 4, whereas 50 µM AP5 completely abolishes the training effect so that trained and untrained trajectories overlap. **C)** End-of-week (average of days 2-4) comparisons consolidate these findings. Trained cultures maintained in control medium or supplemented with BDNF+ show significantly higher normalized spike counts (mean ± SEM) than those treated with AP5, while the AP5 untrained group unexpectedly exceeds its control counterpart. **D)** End-of-week comparisons of unique neurons (mean ± SEM) identified by spike sorting. The number of unique neurons significantly rises with training in control and BDNF+ media but remains low under NMDA-receptor blockade, consistent with AP5 limiting activity-dependent neuronal recruitment. These results demonstrate a robust, stimulation-driven LTP in CNS-3D organoid networks that can be amplified by exogenous BDNF and suppressed by NMDA-receptor antagonism, establishing the platform’s utility for pharmacological interrogation of synaptic plasticity.

**Table 1.** Mixed-effects ANOVA and post hoc comparisons of LTP by dosing group. The upper section shows omnibus mixed-effects ANOVA results for the effects of training and training-by-time interaction within each dosing group. The lower section shows post hoc comparisons between trained and untrained groups at each testing day, reported as estimated mean differences with standard errors and p values.

| Dosing Group | Factor Comparison | df | F Value | P Value | Residual SD |
| --- | --- | --- | --- | --- | --- |
| Control | Training | 1:22 | 10.69 | 0.0035 | 67.83 |
|  | Training × Time | 4:88 | 1.77 | 0.14 | 67.83 |
| BDNF+ | Training | 1:22 | 48.04 | 5.85E-07 | 154.69 |
|  | Training × Time | 4:88 | 3.72 | 0.0076 | 154.69 |
| AP5 | Training | 1:22 | 0.68 | 0.42 | 71.85 |
|  | Training × Time | 4:88 | 0.27 | 0.89 | 71.85 |

Post-hoc trained vs untrained comparisons
| Dosing Group | Testing Day | df | Estimate | SE | P Value |
| --- | --- | --- | --- | --- | --- |
| Control | Day 1 | 76.29 | 31.83 | 34.37 | 0.36 |
|  | Day 2 | 76.29 | 89.25 | 34.37 | 0.011 |
|  | Day 3 | 76.29 | 86.78 | 34.37 | 0.014 |
|  | Day 4 | 76.29 | 126.91 | 34.37 | 4.20E-04 |
|  | Day 7 | 76.29 | 48.76 | 34.37 | 0.16 |
| BDNF+ | Day 1 | 102.66 | 141.11 | 70.07 | 0.047 |
|  | Day 2 | 102.66 | 203.01 | 70.07 | 0.0046 |
|  | Day 3 | 102.66 | 363.36 | 70.07 | 1.10E-06 |
|  | Day 4 | 102.66 | 438.84 | 70.07 | 8.90E-09 |
|  | Day 7 | 102.66 | 199.16 | 70.07 | 0.0054 |
| AP5 | Day 1 | 63.21 | -2.15 | 38.86 | 0.96 |
|  | Day 2 | 63.21 | -42.97 | 38.86 | 0.27 |
|  | Day 3 | 63.21 | -20.67 | 38.86 | 0.60 |
|  | Day 4 | 63.21 | -32.77 | 38.86 | 0.40 |
|  | Day 7 | 63.21 | -20.04 | 38.86 | 0.61 |

To assess the stability of this potentiation, we continued evaluation after a two-day pause with no stimulation. In the control condition, trained cultures showed a partial reduction in spike output by Day 7, indicating that potentiation was not fully maintained without ongoing stimulation. This decay pattern is characteristic of early-phase LTP (E-LTP), which is known to last hours to days and typically decays in the absence of protein synthesis or further reinforcement. By contrast, in the BDNF-supplemented condition, trained cultures showed a decrease in activity but still maintained significantly elevated spike counts after the weekend pause, suggesting that BDNF promoted the stabilization and persistence of potentiation. This prolonged effect is consistent with a transition toward late-phase LTP (L-LTP), which involves long-term structural or molecular changes that are dependent on neurotrophic signaling pathways.

We compared end-of-week metrics across conditions (Fig. 4C) by averaging the normalized spike counts from days 2-4 (Wednesday-Friday) of the week. The average normalized spike count for trained organoids in control and BDNF+ conditions were found to be significantly higher than that in AP5-treated organoids. Similarly, the number of active neurons detected via spike sorting were examined (Fig. 4D) comparing trained vs untrained samples for each dosing condition. In both control and BDNF groups, the trained organoids exhibited a larger count of distinct neurons contributing spikes by the end of training compared to untrained, whereas under AP5 the trained and untrained had equally low counts.

### Organoids trained in a closed-loop maze game

Since patterned stimulation alone could induce plasticity, we next evaluated whether the activity of organoid pairs could be modified as learned responses to reward (rest) or penalty (high-frequency stimulation). We designed a neural interface for the organoids with a custom-designed maze game, inspired by Pac-Man®, under a closed-loop protocol broken into learning and memory phases (Fig. 5A), effectively treating the organoid pair as a living agent navigating a simple environment. In this setup, the organoid could only influence the game through its neural responses to stimuli, which the controller translated into directional movements, and in turn the organoid received sensory consequences in the form of stimuli representing “reward” or “penalty”. This embodied closed-loop format is conceptually similar to recent studies where neural cultures learned to play games when given feedback^52,53^.

**Figure 5.**
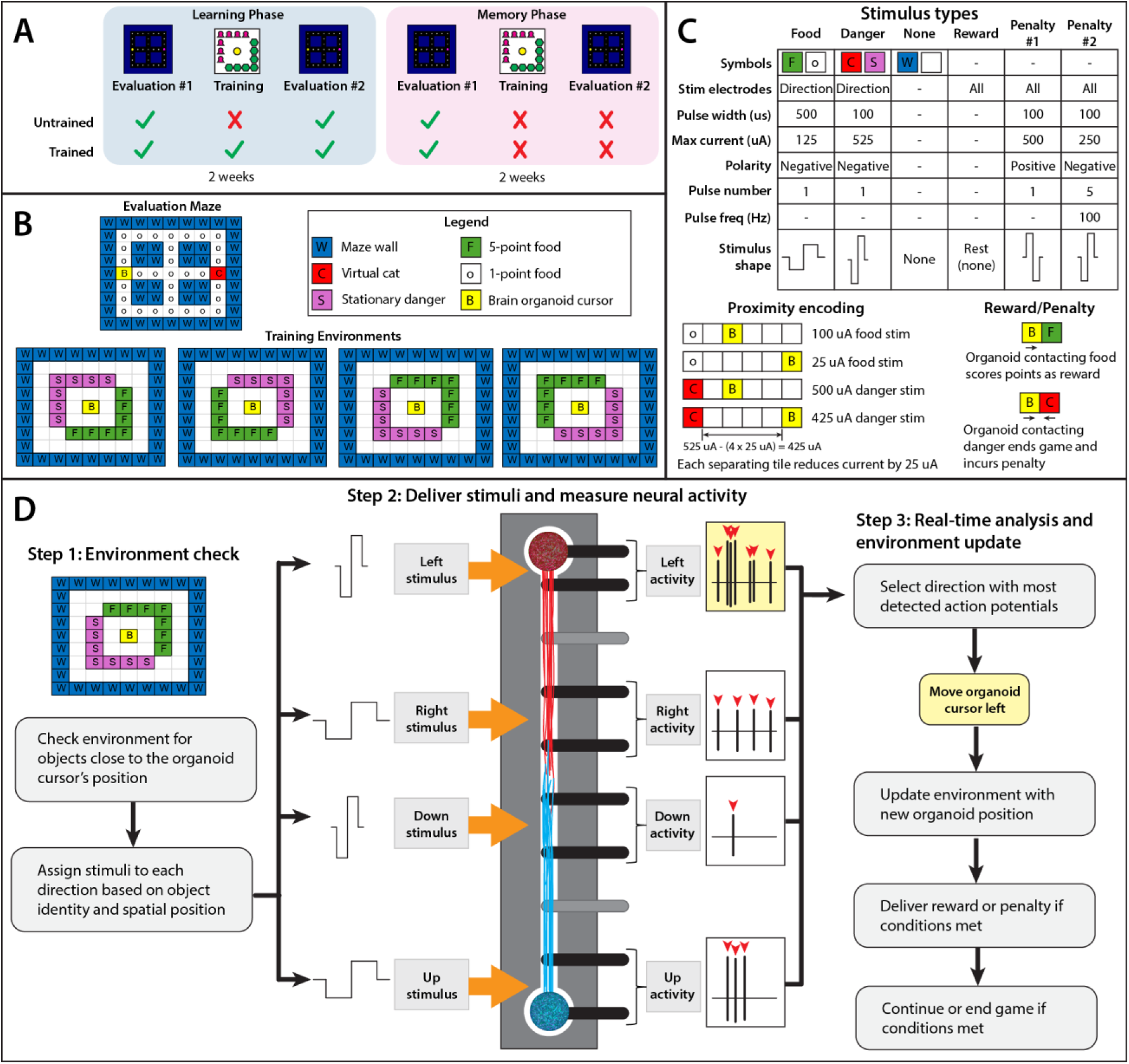
Organoid responses to stimuli associated with direction of movement and proximity of reward/penalty in a maze-game task. **A)** Experimental schedule. Evaluation #1 (pre-training) consists of three closed-loop maze runs for both untrained and trained cohorts (maximum 60 s per run). The trained cohort then completes a training block: six trials in each of four training mazes (24 total runs, 15 s maximum each). These short mazes repeatedly force the cursor to choose between a food tile (reward stimulus) and a danger tile (penalty stimulus); failure to contact either within 15 s also yields the penalty. The untrained cohort rests during this interval. Finally, Evaluation #2 repeats three runs of the evaluation maze for both cohorts under identical closed-loop conditions, yielding six evaluation trials per cohort in total. **B)** Maze layouts. The single evaluation maze (top left) is used in both Evaluation #1 and #2. Four training mazes (bottom) are rotated permutations of the same geometry to prevent directional bias. Legend (right) defines tile types used in maze layouts. **C)** Stimulus library. For each tile category - Food, Danger, Background (None), Reward, Penalty-1, Penalty-2 - the accompanying table specifies: addressed electrode pair, biphasic-pulse width, maximum current, polarity, pulse count, pulse frequency, and a schematic waveform. Proximity (lower left) is encoded by amplitude: every grid step between the cursor and a stimulus source reduces stimulus current by 25 µA. The lower right inset illustrates the conditions for scoring a reward versus incurring a penalty. **D)** Closed-loop control cycle. **Step 1:** the controller polls tiles in orthogonal directions (left, right, down, up) and assigns stimulus parameters according to object identity and distance. **Step 2:** biphasic trains are applied simultaneously to the two electrodes that define each direction while multi-unit activity is recorded. **Step 3:** real-time spike counting selects the direction with the highest firing rate, moves the cursor accordingly, updates the virtual environment, and triggers reward or penalty stimuli when conditions are met. A trial terminates on cursor–danger contact or at the time-out limit (training trials also end on food capture). This framework enables direct comparison of learning efficiency, memory retention, and stimulation-dependent modulation of long-term potentiation (LTP) and depression (LTD) between trained and untrained brain-organoid networks.

Figure 5B outlines the maze layouts used. During evaluation trials (before and after training), a fixed evaluation maze was used for all organoids, containing “food” and a single pursuer (virtual cat) at specific locations. During training trials, four smaller mazes were used (Fig. 5B, bottom), each being a rotated version of simple binary decisions (two directions leading to food, the other two to danger). This prevented any directional bias as each cardinal direction had an equal chance to be correct across trials. The training mazes also forced the organoids to experience both rewarding and penalizing outcomes. The legend in Fig. 5B defines the tile types and their associated stimuli. We crafted a library of stimuli (Fig. 5C) such that each tile or event corresponded to a unique electrical input pattern. For instance, when the cursor was adjacent to the food tile, a specific “food proximity” stimulus was delivered on electrodes corresponding to the direction of the food. Contact with the food tile triggered a reward stimulus, represented by a rest period without stimulation intended to reinforce whatever neural activity led to that success. Conversely, approaching the danger tiles elicited a “danger proximity” stimulus, and moving onto it triggered one of two penalty stimuli (see below for details) as a form of negative reinforcement. Importantly, the intensity (i.e. current amplitude) of these stimuli was attenuated with distance: for every grid step farther from a tile, the stimulus current was reduced by 25 µA per step. Thus, the system was designed so that the organoid network would receive a gradient of stimulus strength indicating how close it was to a goal or hazard, an element akin to sensing in a real environment. All stimuli were delivered as biphasic pulses or pulse trains on designated electrode pairs, targeting both organoids and/or the channel as appropriate to engage a broad network response (Fig. 5C table lists the stimulus types and parameters).

The closed-loop control cycle (Fig. 5D) ran continuously during a maze or training trial. In each step, the system applied four directional stimuli simultaneously (up, down, left, right) and listened to the organoid activity for ∼0.1 s. Because these stimuli were spatially separated on different electrode pairs, the organoid’s response could exhibit bias toward one direction (e.g. higher firing on electrodes associated with “up”). The controller then moved the cursor in the direction that elicited the highest neural firing rate using the stimulus artifact removal and spike identification methodology used in the LTP studies. This effectively meant that the organoid “chose” the direction by generating more activity in response to that directional cue. We note that initially, these choices may be random or reflect innate biases in connectivity (e.g., one organoid might be more responsive to stimuli on a certain side or show a preference to move towards danger), but with training, the goal was to see if the organoid could learn to favor the direction leading to reward (food) over that leading to penalty (danger). After the move, if a reward or penalty condition was met, the corresponding stimulus was delivered, which could further alter the network state, and if the trial ended (by reaching food or danger or timeout), the next trial would start after a 30 second inter-trial interval. Each organoid pair’s performance was tracked over multiple trials to assess learning in the form of the points scored for contacting food or survival time in game iterations, both measured as best across three repetitions.

Using this closed-loop framework, we compared a trained group, which underwent the full maze training regimen, to an untrained group that only did the evaluation runs without any intervening training. Additionally, within the trained group, two different penalty feedback strategies were tested: Penalty #1 was a single, inverted polarity danger stimulus and Penalty #2 was a more intense burst of 5 high-frequency danger-like pulses (Fig. 5C). The rationale was to see if a stronger punishment signal (Penalty #2, involving high-frequency stimulation known to drive plasticity) would be more effective for learning than a weaker one (Penalty #1) that seemed aversive due to lack of responses (Fig. 3F). Both paradigms used the same reward stimulus for consistency.

### Organoid learning of maze-game task was dependent on BDNF

Across two weeks of closed-loop training, we found that organoid pairs receiving the Penalty #2 paradigm (high-frequency burst punishment) showed significant signs of performance improvement, whereas those receiving the Penalty #1 paradigm did not. Figure 6A plots the raw and normalized point totals for three groups – untrained (N = 8 organoid pairs), trained with Penalty #1 (N = 15 organoid pairs), and trained with Penalty #2 (N = 10 organoid pairs) – over the average learning and memory weeks. Upon initial observation, the raw performance (Fig. 6A left; best points scored across 3 maze runs) of the groups show substantial overlap but the scores for Penalty #2 became significantly elevated compared to untrained by the end of the week. When normalized to each group’s Day 0 baseline (Fig. 6A, middle), a trend appeared as early as Day 2 where the Penalty #2 group demonstrated a significant, monotonic increase over time, whereas the other groups showed little or no upward trend. This was corroborated by examining the average performance metrics (Fig. 6A right) at the end-of-week by averaging the normalized values for each condition across days 2-4. The end-of-week analysis confirmed that only Penalty #2 induced significant improvement in learning, a benefit that was lost in the memory phase without continued reinforcement. Survival time outcomes mirrored these findings (Figure 6B), with Penalty #2 extending survival progressively during learning, while Penalty #1 provided a smaller, but significant, benefit. No group differences were observed in the memory phase, highlighting the transient nature of the learning effect.

**Figure 6.**
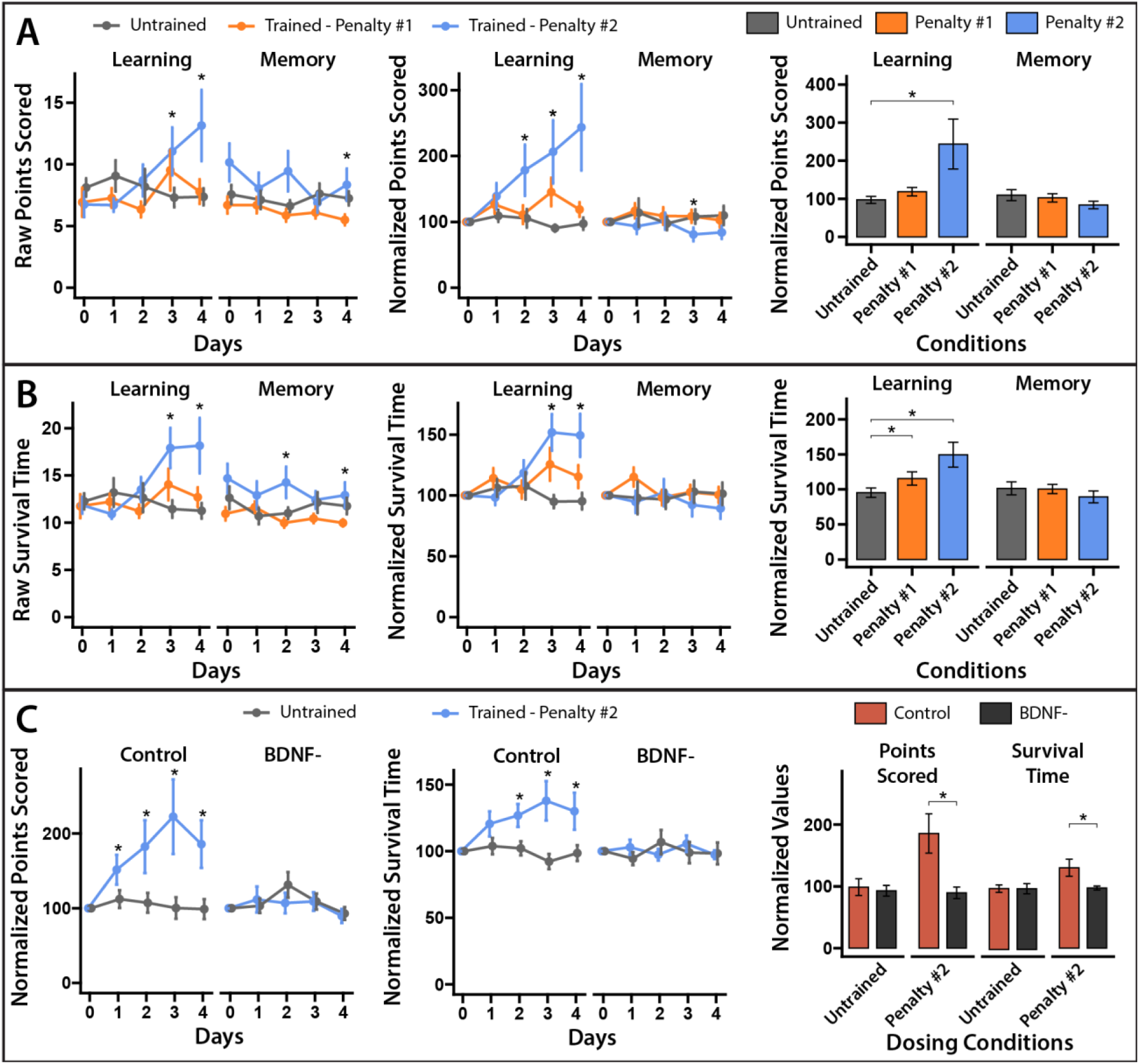
Organoid learning of closed-loop maze-game task depended on high-frequency penalty stimulus and the presence of BDNF. **A)** Best-score analysis for the three consecutive maze runs performed each day during Evaluation #1. The left sub-panel plots raw points (mean ± SEM) accumulated during the average learning and memory weeks, revealing overlapping data across groups though Penalty #2 still outperforms the untrained condition. When each sample is normalized to its Day 0 performance (middle sub-panel; mean ± SEM), a clear upward trajectory emerges for cultures trained with Penalty #2 (five-pulse, 100 Hz danger burst) during the average learning week, whereas Penalty #1 (inverted-polarity danger stimulus) remains indistinguishable from untrained controls. The right sub-panel collapses Days 2–4 into an end-of-week mean ± SEM, confirming that only Penalty #2 produces a significant improvement during learning, an effect that vanishes without reinforcement in the memory phase. **B)** Survival-time outcomes follow a similar pattern. Raw values and their normalized counterparts both show that Penalty #2 lengthens survival progressively during learning; Penalty #1 confers a smaller but still significant benefit, while no group differences persist in memory. Averaged end-of-week data reiterate the superiority of Penalty #2 for extending survival time. **C)** Maze performance depends on exogenous BDNF. In a replication run conducted in control medium containing 20 ng mL⁻¹ BDNF, training with Penalty #2 again elevates normalized points and survival time (mean ± SEM) relative to untrained wells. Removing BDNF from the medium abolishes these training-related gains, as shown both in the day-by-day trajectories and in the pooled end-of-week comparisons, indicating that the behavioral advantage conferred by high-frequency penalty bursts requires BDNF signaling. Together, these data demonstrate that a high-frequency burst penalty is more effective than an inverted-polarity penalty at enhancing goal-directed performance and persistence, and that this enhancement is contingent on the presence of BDNF, mirroring the pharmacological modulation observed for long-term potentiation in open-loop experiments.

Next, we compared organoid learning under normal control media versus BDNF-depleted conditions to determine the possibility for pharmacological modulation. As shown in Figure 6C, organoids trained with Penalty #2 in our standard medium (supplemented with 20 ng/mL BDNF) displayed a significant improvement in both total points (left) and survival time (middle) relative to untrained controls. However, when the same training protocol was applied to organoids in BDNF-free medium, the trained networks no longer outperformed the untrained with points and survival time equivalent to untrained controls. This is shown directly in the right panel of Figure 6C, where the control medium condition is compared to a BDNF-condition by averaging data across days 2-4 (end-of-week).

### Transcriptomic changes mirror functional changes with training and dosing

We profiled bulk RNA-seq (N = 3 samples per condition, each composed of 2 organoid pairs) across open-loop LTP and closed-loop maze-game cohorts to test whether training state, pharmacology, and time map onto gene expression for learning, memory, and synaptic plasticity. The study design (Fig. 7A) spanned three comparisons: dosing at DIV17 in LTP (trained versus untrained in Control, BDNF+ 50 ng mL⁻¹, and AP5), time in LTP Control (trained and untrained at DIV17 versus DIV24), and maze-game at DIV46 (trained versus untrained in Control versus BDNF−). UMAP on all normalized gene counts separated maze-game from LTP samples, consistent with differences in age or batch-to-batch organoids (Fig. 7B, left). In contrast, UMAP on GSVA scores derived from learning, memory, and synaptic plasticity gene sets organized samples along a functional axis: trained Control and trained BDNF+ at DIV17 in LTP, together with trained Control maze-game, clustered in the upper right, while AP5 (trained or untrained), LTP Control at DIV24, and maze-game BDNF− clustered in the lower left (Fig. 7B, right).

**Figure 7.**
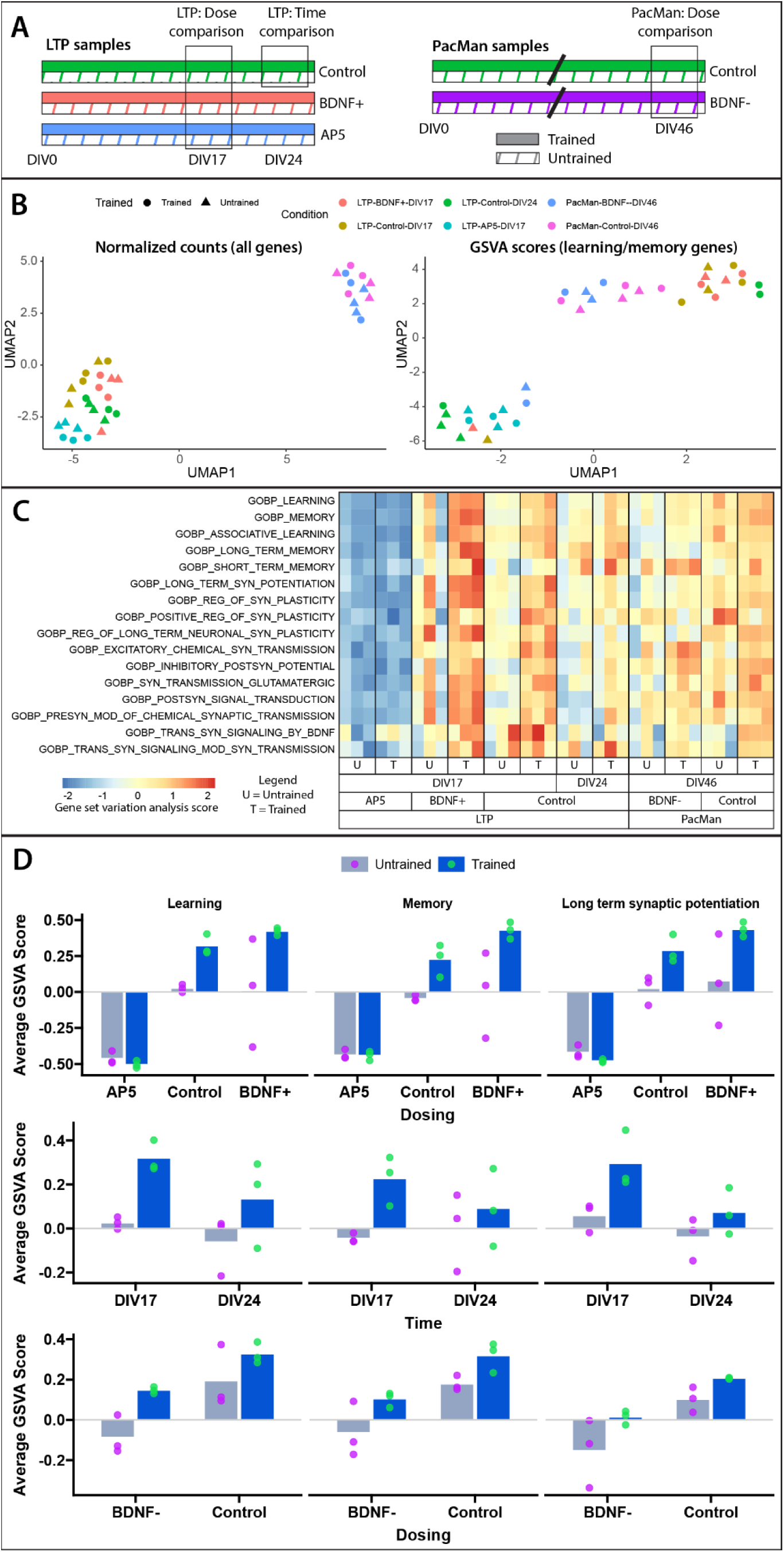
RNA-seq reveals condition- and training-dependent transcriptional programs that aligned with pathways associated with synaptic plasticity. **A)** Schematic of experimental timelines and dosing conditions for the LTP (DIV17, DIV24) and maze-game (DIV46) cohorts. Black boxes denote the three bulk RNA-seq comparisons: LTP dosing at DIV17 (trained versus untrained in Control, BDNF+ [50 ng mL⁻¹], and AP5), LTP timepoint in Control (DIV17 versus DIV24), and maze-game at DIV46 (trained versus untrained in Control and BDNF− medium). **B)** UMAP projections computed from all normalized gene counts (left) or from GSVA scores for learning-, memory-, and synaptic plasticity-related gene sets (right). **C)** Heat map of GSVA scores across 15 GO terms associated with learning, memory, LTP, and synaptic plasticity (n = 3 samples per group). **D)** GSVA summary scores displayed as bar plots with individual samples overlaid; rows correspond to dosing (top), LTP timepoint (middle), and maze-game medium condition (bottom), and columns indicate Learning, Memory, and Long-term Synaptic Plasticity.

Gene set variation analysis reinforced these patterns. Heat maps across 15 GO sets showed uniformly low scores in AP5 irrespective of training, higher scores in trained BDNF+ and trained Control at DIV17 relative to their untrained counterparts, and a decline by DIV24 in LTP Control that paralleled the cessation of training (Fig. 7C). Maze-game samples displayed training-associated increases under Control medium, whereas BDNF− remained comparatively low. Quantitative summaries confirmed these trends across learning, memory, and long-term synaptic plasticity: in the dosing comparison at DIV17, trained groups exceeded untrained, AP5 was depressed, and trained BDNF+ surpassed trained Control with the largest margin in the memory group; in the time comparison, trained Control scores decreased from DIV17 to DIV24 while untrained remained relatively stable; in maze-game, Control exceeded BDNF− for both training states, and trained Control exceeded untrained Control across metrics (Fig. 7D).

## Discussion

This study clearly demonstrated that a structured, human brain organoid network can express key hallmarks of learning-related plasticity, including network-level long-term potentiation (LTP) and adaptive responses in a closed-loop game task. By integrating paired CNS-3D organoids with guided axonal connectivity on an embedded electrode array (EEA), we created a platform in which distributed neural activity can be stimulated, recorded, and modulated over time. Importantly, these findings move brain organoids beyond models of spontaneous activity alone toward functional in vitro systems that can probe mechanisms of human learning and memory. In the context of CNS drug discovery, where current biomarkers and preclinical models often fail to predict human efficacy, this platform suggests a path toward more human-relevant functional readouts of neural plasticity.

A key advancement of this study is the demonstration of long-term potentiation (LTP) in a multi-organoid network coupled through axonal tracts and validated by pharmacological perturbation. Potentiation that persisted across days was abolished by NMDA receptor blockade with AP5 and enhanced by BDNF supplementation, consistent with canonical NMDA receptor-dependent, neurotrophin-facilitated plasticity and indicating that these interconnected organoid circuits support durable synaptic strengthening in vitro^9,15,54,55^. Transcriptomic profiling further reinforced this interpretation, as GSVA-based analyses of learning, memory, and synaptic plasticity gene sets tracked functional performance across both LTP and maze-game cohorts, with trained Control and BDNF+ conditions showing stronger enrichment than untrained or AP5-treated samples. Together, these findings indicate that the EEA-connectoid system captures biologically meaningful and pharmacologically tunable forms of plasticity rather than nonspecific stimulation effects.

The closed-loop experiments further suggest that these organoid networks used patterned feedback to modify responses in a virtual environment. Organoids exposed to appropriately strong feedback improved navigation toward reward, indicating a form of adaptive, goal-directed learning. At the same time, this effect was transient and dependent on exogenous BDNF support, with performance declining in its absence. These observations support the existence of feedback-dependent, short-lived functional adaptation in human organoid networks. The interpretation is significant because it advances organoids from passive electrophysiological preparations toward systems in which plasticity and learning can be evaluated through task performance. Compared with prior closed-loop studies in 2D neural cultures, such as the Pong-like game in Kagan, et al ^53^, this 3D paired-organoid system provides a more physiologically relevant human neural architecture by incorporating richer synaptic connectivity, neuron-glia interactions, and guided long-range axonal connectivity, enabling adaptive game performance to be evaluated alongside network-level plasticity and pharmacologic modulation. These features of the 3D system may help explain the larger improvement in game performance observed here, as well as the longer-lasting retention of training effects over hours to days compared with the more modest score changes and shorter timescales reported in prior 2D work^53^. More broadly, these findings represent a step toward organoid intelligence, in which human neural tissues are studied not only as models but also as living computational systems capable of plasticity-dependent, task-relevant adaptation^42–46^. This capability may be especially useful for translational applications, including screening compounds that enhance or impair synaptic plasticity, testing patient-derived disease organoids for deficits in learning-related network responses, and testing therapies designed to mitigate disease-related cognitive dysfunction^56,57^.

Future work will focus on several opportunities to further strengthen this platform. Variability in electrode coupling remains a challenge that will need to be reduced for robust translational use. In addition, the memory-like effects observed here were not stably maintained without reinforcement, suggesting that the present system more reliably models inducible potentiation and organoid learning rather than persistent long-term memory. More durable responses may require longer training paradigms, greater cellular maturity, added myelinating or glial support, or stimulation patterns that better mimic consolidation-related processes^58^. The current network scale, on the order of ∼25,000 neurons per organoid^59^, is also sufficient for simple assays but will likely need to expand for more complex computations or behaviors. Finally, as organoid systems become more functionally sophisticated, ethical oversight must evolve in parallel^60,61^. Overall, our findings establish a human organoid platform that combines engineered connectivity, chronic electrophysiology, and closed-loop feedback to model core features of learning-related plasticity, providing a promising foundation for mechanistic neuroscience, CNS drug discovery, and future studies of computation in living neural tissue.

## Acknowledgements

The authors thank Dave Weiner, Chris Butt, and Alif Saleh (28bio) for their contributions developing and conceptualizing both the open-loop and closed-loop implementations of this work and especially guiding the future applications. The authors would also like to thank Thomas Hartung and Lena Smirnova (Johns Hopkins University) for feedback on experimental design, results, and future directions. We also thank Alysson Muotri and his lab for the use of the CNS-3D organoid license used in this work.

## Methods

### Coating Procedure

For Matrigel coating, an aliquot of Matrigel (Corning, 354277) was placed on ice until mostly thawed, allowing a small piece of ice to remain. Meanwhile, 25 mL of cold DMEM/F12 (50/50) (Corning, 10-092-CV) media was aliquoted into a 50 mL conical tube. A P1000 pipette tip was cooled by aspirating the cold media several times, then approximately 500 µL of the media was left in the tip and added to the Matrigel aliquot. The mixture was gently mixed in the Eppendorf tube to ensure complete thawing, then transferred to the conical containing the cold media. The Eppendorf tube was rinsed twice with portions of the cold media to recover all of the Matrigel. The final coating solution was mixed thoroughly using a serological pipette.

Subsequently, 2.5 mL of the matrix solution was added to each 60mm cell culture dish (Cell Treat, 229660), sufficient to coat ten plates. Plates were gently swirled to ensure even coating and then incubated at 37°C for at least one hour or preferably overnight. Plates older than four days were avoided due to potential compromised cell attachment. Prior to use, unbound material was aspirated, and cell suspension or media was added immediately without washing to prevent the surface from drying.

For Laminin coating using Cultrex® Mouse Laminin I (Biotechne, 3400-010-02), the product was thawed at 2–8°C and kept on ice when ready. It was diluted 1:100 in PBS-Ca-Mg (Corning, 21-040-CV) to a concentration of approximately 10 µg/mL and gently homogenized by pipetting. The appropriate volume of the diluted solution was added to each well of the tissue culture plate, ensuring complete coverage of the well bottom. Plates were then incubated at 37°C for four hours or overnight as recommended. When ready for use, the Laminin solution was aspirated, the wells were washed once with PBS-Ca-Mg, and cell suspension was added immediately without allowing the coated surface to dry.

### Media Preparation

mTeSRT™1(Stem Cell Technologies, 85850) was prepared by following Stemcell Technologies’ Technical Manual Maintenance of Human Pluripotent Stem Cells. Use sterile techniques to prepare complete mTeSR medium (Basal Medium + 5X Supplement). The following example is for preparing 500 mL of complete medium. The 5X Supplement was thawed to room temperature and mixed thoroughly. If the supplement appeared slightly cloudy, a rare but possible occurrence, it was briefly placed in a 37°C water bath and swirled until clear, for no longer than five minutes. Once fully thawed and clear, 100 mL of the supplement was added to 400 mL of mTeSR Basal Medium and mixed thoroughly without filtering, to avoid loss of essential macromolecules. If not used immediately, the complete medium was stored at 2-8°C for up to two weeks, or aliquoted (40 mL into 50 mL conical tubes) and frozen at -20°C for up to six months. Upon thawing, the medium was used immediately or stored at 2-8°C for up to two weeks, but was never refrozen.

To prepare Neural Basal Complete medium (NBC), 0.5x N2 supplement (Thermo Fischer, 17502-048), 0.5x B27 supplement (Thermo Fischer, 17504-044), and 1x Penicillin-Streptomycin (Hyclone, 16777-164) were added to 500 mL of DMEM/F12 (50/50) containing Glutamine and Hepes. The resulting medium was stable at 4°C for up to one month and protected from light. Aliquots (40 mL) could be frozen in 50 mL conical tubes and stored at -20°C for up to six months. For NBF, a fresh aliquot of bFGF (Thermo Fisher, PHG0023) was thawed on ice and briefly centrifuged. Then, 8 µL of bFGF was added to 40 mL of pre-warmed NB medium (yielding a final concentration of 20 ng/mL) immediately before applying the medium to cells.

To prepare Neural Derivation Medium (NDM), 10 mL of B27 supplement without vitamin A (Thermo Fisher PHG0313), 1x Penicillin-Streptomycin, and 1x Glutamax (Gibco, 35050-061) were added to 500 mL of Neurobasal medium. This medium was also stable at 4°C for one month if shielded from light and could be aliquoted (40 mL per 50 mL tube) and stored at -20°C for up to six months. For NDEF, fresh bFGF and EGF aliquots were thawed on ice, centrifuged briefly, and 8 µL of each was added to 40 mL of pre-warmed ND medium (achieving 20 ng/mL of each factor) before applying it to cells.

### Induced Pluripotent Stem Cell (iPSC) Differentiation

iPSCs were differentiated into NPCs as described previously^62,63^. iPSCs colonies were lifted off and kept in suspension, under rotation (95 rpm) for 7 days to form embryoid bodies (EB) in DMEM/F12 50:50 with 1x Glutamax), 1x N2, 1x penicillin-streptomycin, 10-μm SB431542, and 1-μm dorsomorphin.. EBs were gently disrupted, plated onto Matrigel-coated plates, and cultured in DMEM/F12 50:50 with 1x HEPES, 1x Glutamax, 1x PS, 0.5x N2, 0.5x Gem21 NeuroPlex (Gem21; Gemini Bio-products), supplemented with 20-ng/mL basic fibroblast growth factor (bFGF; Life Technologies) and 20ng/mL of epidermal growth factor (EGF; Life Technologies) for 7 days to generate neural rosettes. Neural Rosettes were then manually isolated and enzymatically dissociated into a single-cell suspension. These p1 Neural Progenitor Cells (NPCs) were then expanded up t to create a large working bank for the CNS-3D organoids.

### CNS-3D organoid generation from NPCs in 384 well format 3D

NPC’s are thawed on a monthly basis to create regular builds of CNS-3D organoids. NPC’s are expanded 3 times to then seeded into 384-well low-attachment plates using a BioTek Liquid Handling System, then centrifuged and transferred to a 37°C incubator for culture.

### CNS-3D Organoid Maintenance

BrainPhys Complete Media (BPC) was prepared using BrainPhys™ Neuronal Medium with SM1 Kit (StemCell Technologies, 05792) containing 1× penicillin-streptomycin and supplemented with recombinant human brain-derived neurotrophic factor (BDNF) and recombinant human glial cell line-derived neuro-trophic factor (GDNF) (StemCell Technologies; BDNF: Cat# 78005; GDNF: Cat# 78058), each at a final concentration of 20 ng/mL. Each Monday, Wednesday, and Friday organoid cultures received a half media change with the BioTek Liquid Handling System before being placed in the custom, pre coated embedded electrode array (EEA) plate. Organoids were 2, 3, and 5 weeks old before being placed in the EEA plates while optimizing this assay.

### EEA Culture Preparation

Custom EEA plates were sterilized and coated with 0.01% Polyethyleneimine (PEI) (Sigma-Aldrich, 181978) in 1X Borate Buffer (Thermo Fisher, 28341) for 1 hour at 37°C temperature. After the incubation period the plates dried for 8 hours and were subsequently coated with 10µg/mL Laminin solution in PBS (Caisson Labs, PBL06) and incubated for minimum 8 hours at 4°C temperature. After coating, the laminin solution was removed. Before seeding the coculture organoids, the plates are equilibrated with 200µL per well of BPC culture media in the 37°C in the incubator.

### EEA Culture

Individual organoids were initially transferred from the U-bottom plate into a sterile 100mm petri dish under the BSL2 safety cabinet. The petri dish and lid was transferred to the clean bench where the equilibrated EEA plate was stored. The warmed media was removed, and the organoids were transferred with micro-forceps from the petri dish to the top bulb of the EEA channel and either the 3^rd^ or 4^th^ bulb (2mm and 4mm down the channel) using micro-forceps (Dumostar, 11294) with a microscope in a Thermo Fisher-Scientific Heraguard Eco Clean bench. Eventually, 4mm was the decided positioning. The organoids were covered with 200 µL of BPC media and returned to the incubator. The organoids adhered for 3 days undisturbed in the incubator; on day 3 the entirety of the channel was covered in Matrigel (Corning, 354230) to allow for 3D growth over the electrodes. The matrigel was placed by removing the cultures from the incubator and placing it in the clean bench. Culture media was removed with a pipette to expose the channel. 2-3µL of Matrigel was dispensed into the channel, filling and covering the entirety of the channel and bulbs. After 2-3 minutes, once the matrigel was hardened 200µL of fresh BPC media was dispensed into each well. After this matrigel procedure and media change, a regular Monday, Wednesday, Friday full 200µL media change was established for the following 2-7 weeks. Weekly imaging was performed during the growth weeks to ensure PNS-3D organoid cultures were healthy and robust before dosing took place.

### Imaging methods

Organoid images were taken on the Molecular Devices ImageXpress Micro Confocal Microscope on DIV 4 of the organoid formation phase before organoids were placed in the bulb of the EEA channel to ensure all organoids had formed correctly. The TL25 images were taken with a 4X objective with a 0.05ms exposure time. Weekly imaging was performed once per week to ensure the health and growth of the cultures. Cultures were imaged with the Molecular Devices ImageXpress Micro Confocal Microscope in TL25, with a 10X objective for 7.27ms exposure. Z planes of 19 steps across 8 sites down the channel allowed us to visualize the 3D nature of the peripheral nerve. The range was 15µm between steps for a total range of 270µm. During the dosing week, cell cultures were imaged 2 additional times, once in the middle of the week and once at the end of the seven-day dose period. Images were stitched together with a custom Matlab application for growth analysis and monitoring.

### Dosing

After two weeks of EEA culture, organoids underwent 2 weeks of compound testing for the remainder of culture. For the LTP study cultures underwent full media changes every Monday, Wednesday and Friday with 200µM AP5 (Medchem Express, HY-100714) in Brainphys Complete Media, 50ng/mL BDNF in Brainphys Complete Media, and control media. For the maze study, 50ng/mL BDNF in Brainphys Complete Media, Brainphys Complete media without BDNF, and control media were applied in the same manner but for four weeks while undergoing electrophysiological training and assessment.

### Stimulation and Recording Methods

All electrophysiology experiments were conducted using a custom 24-well embedded electrode array (EEA) plate that allows concurrent stimulation and recording from paired organoids. Each well contained ten planar microelectrodes (50 µm diameter) embedded in the floor of the inter-organoid channel, along with integrated reference electrodes at each end of the channel (1.2 mm diameter) to provide a common ground. Custom headstages incorporating Intan RHS2116 stimulator/amplifier chips (Intan Technologies) were used to interface the EEA plate with the recording hardware. Two data acquisition systems were utilized: an Intan RHS Stim/Recording Controller running RHX v3.1.0 software, and an Alpha Omega AlphaRS Pro system running AlphaRS proprietary software. The Intan RHS system is a 128-channel-count system interfaced with a custom headstage allowing 12 of 24 wells to be stimulated/recorded simultaneously in open-loop configuration. The AlphaRS Pro is a 256-channel-count system capable of simultaneous stimulation/recording of all 24 wells using a custom headstage in both open-loop and real-time closed-loop control. Signals from the RHS2116-based headstages were digitized at 30 kHz and band-pass filtered (analog passband 80–7500 Hz) on-board to capture action potentials. During stimulation, the Intan amplifiers engaged an artifact suppression mode by temporarily reducing the high-pass cutoff to ∼1 kHz to minimize stimulus artifacts. Recorded data were saved in Intan’s RHX format (.rhd) or a MATLAB-based format on the AlphaRS system. All stimulation used current-controlled, biphasic pulses with phase durations matched and opposite in polarity to ensure charge balance. Each electrode could serve for both recording and stimulation, with the stimulus waveform routed through the RHS2116 stimulator channels under software control. The stimulation and recording protocols were implemented in two modes: open-loop and closed-loop (Figure 1C). In open-loop experiments, predefined stimulation patterns were delivered to the organoids while neural responses were recorded, without any real-time feedback. In closed-loop experiments, stimulation parameters were dynamically adjusted based on the organoid network’s ongoing activity (see Closed-Loop Maze Paradigm below).

### Open-Loop Stimulation

Open-loop stimulation protocols were designed to probe and induce synaptic plasticity. We systematically explored a range of stimulus parameters to optimize evoked responses. This included varying the pulse amplitude (e.g. 25–400 µA), pulse width (100–500 µs per phase), number of pulses in a train (single pulse vs. short high-frequency bursts of 5 or 10 pulses), pulse frequency (e.g. 50 Hz or 100 Hz for bursts), and pulse polarity (standard cathodic-first vs. inverted polarity anodic-first for certain aversive stimuli). Each stimulation “trial” consisted of delivering a specific pulse or pulse train on one or more electrodes and recording the post-stimulus response. A custom MATLAB 2024a application orchestrated all stimulus delivery and data acquisition. Stimulation protocols were defined in a spreadsheet script (Excel format) listing the sequence of stimulus parameters (electrode selection, current amplitude, pulse width, polarity, number of pulses, and trial timing). The MATLAB controller interfaced with the Intan RHX software via TCP/IP or with the AlphaRS Pro API to load each stimulus configuration and trigger synchronized stimulation/recording in the hardware. This setup enabled rapid switching between different stimulus conditions and automated execution of complex stimulation paradigms across the 24-well plate. Parameter tuning experiments (see Results, Fig. 3D) were conducted in open-loop mode to identify stimulus settings that elicited reliable spiking (multi-unit) responses from the organoid networks. Unless otherwise noted, all stimuli were delivered as biphasic pulses with 0.1 ms inter-phase intervals, and trials were separated by at least 1 s to allow network recovery. For plasticity induction (LTP/LTD) studies, repetitive stimulation trains were delivered over several days as described later, with control (unstimulated) organoids maintained in parallel for comparison.

### Closed-Loop Maze Paradigm

For closed-loop experiments, we developed an interactive maze-game paradigm, inspired by Pac-Man®, to provide real-time feedback-driven stimulation. The closed-loop system linked the organoid electrophysiology to a virtual agent navigating a grid-based maze, using principles of reinforcement learning to emulate goal-directed behavior. The game environment was implemented in MATLAB and presented four cardinal directions (up, down, left, right) as possible moves for the agent. In each closed-loop trial, an organoid pair was tasked with guiding the agent from a start location to a target (“food”) while avoiding danger. Custom logic translated neural activity into directional decisions and delivered feedback stimuli based on the agent’s progress.

Each maze was composed of a 2D grid of tiles with designated properties (open path, food, danger, etc.), as illustrated in Fig. 5B. We utilized two types of mazes: evaluation mazes and training mazes. Evaluation mazes (used in pre- and post-training assessments) had a fixed layout with one food target and a computer-controlled pursuer (obstacle) to challenge the organoid without any additional training cues. Training mazes were simpler configurations that required a binary choice (turn toward food vs. toward danger), and we employed four rotational variants of a basic two-choice maze. This ensured that across trials each cardinal direction was equally likely to be “correct,” preventing any directional bias in the organoids’ responses. Organoid pairs underwent daily training sessions in which they experienced a series of these maze trials with feedback, followed later by evaluation trials without feedback to test learned performance. A typical training session consisted of multiple maze runs (e.g. 10–20 trials per day), and training was carried out over 5–7 days (constituting the “learning phase”). In the subsequent week (the “memory phase”), the organoids were tested in evaluation mazes without further feedback to assess retention of any learned behavior. Performance metrics recorded included the total points (rewards) earned per trial and the survival time (time until hitting danger), which were averaged or summed over sets of trials for comparison across days.

The neural interface between the organoids and the maze-game operated on a cycle of stimulate-listen-act that repeated continuously during each trial. At each time step (iteration) of the game, the system delivered four *directional stimuli* simultaneously: one cue for each cardinal direction. These stimuli were delivered through four distinct pairs of electrodes on the EEA (each pair corresponding to one direction), targeting different spatial regions of the organoid network. For example, “up” stimuli might be delivered on a pair of electrodes located under the top organoid, whereas “down” stimuli target electrodes under the bottom organoid, and “left”/“right” stimuli delivered in the connecting channel or opposing sides, such that each direction had a unique spatial stimulation pattern. All directional cues were biphasic pulses with moderate current (outlined in Fig. 5C) delivered concurrently. The stimulation parameters for each cue were optimized to be salient but not evoking overwhelming activity, so that the organoids’ differential response could indicate a “preference.” Immediately after delivering the directional cues, the system entered a listening period (∼100 ms) during which multi-unit firing activity from the organoids was recorded in real time. A fast online analysis computed the firing rate on each of the four directional electrode pairs (using threshold crossing to detect spikes). The direction yielding the highest summed spike rate was interpreted as the organoids’ choice for that step. Consequently, the virtual agent was moved one tile in that chosen direction. This closed-loop decision process effectively allowed the organoid network’s activity to control the agent’s navigation: the organoids “steered” the agent by generating more spikes in response to one of the directional stimuli.

As the agent moved through the maze, context-specific feedback stimuli were provided to reinforce learning. We designed a library of unique electrical stimuli mapped to important game events. For instance, when the agent’s next move would place it adjacent to the food target, a special “food proximity” stimulus was delivered on the electrodes oriented toward the food’s location. This cue consisted of a single cathodic-first, 500 us-100 uA pulse, meant to signal that the goal is near. Likewise, approaching a dangerous tile triggered a “danger proximity” stimulus on the electrodes facing that direction, alerting the organoid of impending penalty. If the agent successfully reached the food tile, a reward stimulus was issued: rather than an electrical pulse, the reward was a 1–2 s pause in stimulation (no stimuli delivered). This lack-of-stimulus period served as a positive reinforcement, under the assumption that whatever neural activity led to the success would be intrinsically strengthened by a brief cessation of perturbation. In contrast, if the agent stepped onto a danger tile, a penalty stimulus was delivered to induce negative reinforcement. We tested two different penalty configurations. Penalty #1 was a single biphasic pulse of inverted polarity (anodic-first) at a relatively high current, intended to be an aversive signal. Penalty #2 was a stronger stimulus: a burst of five high-frequency pulses (e.g. 5 pulses at 100 Hz, cathodic-first) at the 250 uA. Both penalty types were delivered on a broad set of electrodes spanning both organoids and the mid-channel, to engage a large portion of the network. To incorporate spatial information, we modulated stimulus intensity based on the agent’s distance from the target or danger. Specifically, for every grid step further away from the food or danger, the amplitude of the proximity stimuli was linearly decreased by 25 µA (for example, if 75 µA at 1 step away, then 50 µA at 2 steps, 25 µA at 3 steps, etc.). This created a gradient of stimulus strength that the organoids could potentially sense, analogous to sensing a gradient in chemical or sensory cues. Throughout closed-loop operation, the MATLAB control program handled real-time event detection and stimulation with a loop cycle on the order of tens of milliseconds, ensuring low latency feedback. Trial outcomes (success or failure), points earned, and survival time were logged for each run. These performance metrics were later used to quantify learning by comparing the trained organoids’ improvements against untrained controls and across different feedback strategies.

### Data Analysis

Recorded electrophysiological data were analyzed both offline (for open-loop experiments) and online (for closed-loop experiments), using custom scripts in MATLAB and R. Preprocessing of raw voltage traces included artifact removal and spike detection. For open-loop stimulus-response data, each trial’s recording was segmented from –100 ms to +500 ms relative to stimulus onset, using the recorded trigger timestamps. A baseline correction was applied to remove stimulus artifacts: we subtracted a copy of the signal low-pass filtered at 10 Hz (4th-order Butterworth, zero-phase) from the raw trace, effectively eliminating slow drift and residual stimulus offset without distorting fast action potentials. The preprocessed trial matrices were then used to compute quantitative metrics of evoked activity. We detected action potentials (spikes) in the high-pass filtered data using an amplitude threshold set at ∼4–5× the noise standard deviation. Detected spike waveforms were aligned to their trough and then subjected to spike sorting to distinguish putative single neurons. We employed feature-based clustering methods using a wavelet-transform clustering algorithm (adapted from Quian Quiroga et al. 2004) to automatically group spikes into units^51^. Each sorted unit’s spike train was examined across the recording period to ensure stability of waveform shape and firing characteristics. For quality control, units with refractory period violations or unstable waveforms were excluded from single-unit analyses, and the remaining multi-unit activity (including all threshold crossings) was used for overall network activity measures.

### Burst analysis

Burst analysis of spontaneous activity was performed to characterize network oscillations and plasticity. We defined a network burst as a sequence of at least 3 spikes with short inter-spike intervals, using a maximum interval method. Specifically, any spikes occurring with consecutive inter-spike intervals ≤ 170 ms were grouped into a burst, as long as the burst duration exceeded 10 ms and the burst was separated from the next spike by at least 200 ms (minimum inter-burst interval). This algorithm was applied to spike timestamps on each electrode (or each sorted unit), yielding burst onset times and burst durations for each recording. From these, we computed the burst frequency (bursts per minute) and mean spikes per burst for each condition. To visualize changes in bursting over time, we employed kernel density estimation (KDE): burst event timestamps were convolved with a Gaussian kernel (bandwidth on the order of the burst interval) to obtain a continuous estimate of burst rate as a function of time. For example, in assessing the development of connectivity between organoids, we compared the KDE of burst frequency on electrodes under each organoid vs. those in the middle of the channel over the first few weeks of co-culture (see Fig. 3B), which illustrated the emergence of synchronized bursting as axonal connections formed.

### Evoked response analysis

Evoked response analysis for plasticity experiments focused on quantifying changes in spike output due to training. For each organoid pair, we measured the number of spikes (or multi-unit firing rate) in a 0–300 ms window after each stimulus, and in some cases the integrated voltage envelope of the response, to capture subthreshold contributions. Potentiation was evaluated by comparing these metrics before and after the training protocol. For short-term potentiation experiments, we normalized evoked response magnitudes to each sample’s own baseline (pre-training on the same day) value to calculate % change due to training. For long-term potentiation experiments, we instead normalized evoked response magnitudes to each sample’s own baseline at the beginning of each week to calculate % change over time. Similarly, for closed-loop behavioral experiments, performance metrics (total points and survival time) were normalized to each organoid pair’s initial baseline run at the beginning of each week to account for individual differences.

### Statistical analyses

Statistical analyses were carried out to determine the significance of observed changes. Depending on the comparison, we used unpaired two-tailed t-tests for single variable comparisons or repeated-measures ANOVA for multi-day learning trends, with post hoc tests (Tukey’s HSD) for multiple comparisons. Non-parametric alternatives (Wilcoxon signed-rank or rank-sum tests) were used if data violated normality assumptions. Data are generally reported as mean ± standard error of the mean (SEM). A significance threshold of *p* < 0.05 was applied for all tests. To assess pharmacological effects, two-factor analyses were used (treatment × training) to test for interaction effects (e.g., whether NMDA receptor blockade by AP5 influenced the degree of LTP induced, or whether BDNF withdrawal affected learning outcomes). All analyses were implemented in MATLAB R2024a or R (v4.3) using custom-written scripts and built-in statistical libraries.

### Visualization

Visualization of results was done using MATLAB and R (ggplot2), with kernel density plots for firing rate/burst distributions, line and bar graphs for time-course and group comparisons, and scatter plots for single-unit activity metrics. Figures were assembled in Adobe Illustrator, with schematics of the experimental setup created in-house to illustrate the EEA platform and neural interface overview. Each dataset was reviewed for outliers or technical artifacts, and any exclusions are noted in the Results. Together, these data analysis methods enabled us to quantify synaptic potentiation and learning-related changes in the organoid cultures with rigor and reproducibility.

